# Bacteriophage infection produces membrane vesicles in *Escherichia coli* via both explosive cell lysis and membrane blebbing

**DOI:** 10.1101/2020.12.23.424155

**Authors:** Pappu K. Mandal, Giulia Ballerin, Laura M. Nolan, Nicola K. Petty, Cynthia B. Whitchurch

**Author notes:** **Corresponding author**, Cynthia B. Whitchurch.

## Abstract

Membrane Vesicles (MVs) are membrane-bound spherical nanostructures that prevail in all three domains of life. In Gram-negative bacteria, MVs are thought to be produced through blebbing of the outer membrane and are often referred to as outer membrane vesicles (OMVs). We have recently described another mechanism of MV biogenesis in *Pseudomonas aeruginosa* that involves explosive cell lysis events which shatters cellular membranes into fragments that rapidly anneal into MVs. Interestingly, MVs are often observed within preparations of lytic bacteriophage, however the source of these MVs and their association with bacteriophage infection has not been explored. In this study we aimed to determine if MV biogenesis is associated with lytic bacteriophage infection. Live super-resolution microscopy demonstrated that explosive cell lysis of *E. coli* cells infected with either bacteriophage T4 or T7, produced MVs derived from shattered membrane fragments. Infection by either bacteriophage was also associated with the formation of membrane blebs on intact bacteria. TEM revealed multiple classes of MVs within phage lysates, consistent with multiple mechanisms of MV biogenesis. These findings suggest that bacteriophage infection may be a major contributor to the abundance of bacterial MVs in nature.

## Introduction

Membrane vesicles (MVs) are non-replicating, membrane-bound spherical nanostructures that prevail in all the three domains of life (1). Bacterial MVs are involved in diverse biological processes such as pathogenicity (2), horizontal gene transfer (3), biofilm formation (4), and to serve as decoys to defend bacteria from antibiotics (5), antimicrobial peptides (6) and bacteriophage predation (7). Moreover, the abundance of bacterial MVs in coastal and open ocean seawater suggests that these could also have major impact on global biogeochemical cycles (8).

In Gram-negative bacteria MVs are thought to be produced via membrane blebbing which involves protrusion and budding of the outer membrane resulting in the formation of MVs, which are often referred to as outer membrane vesicles (OMVs). Membrane blebbing occurs as a result of cell envelope disturbances caused by either imbalance of peptidoglycan biosynthesis or intercalation of hydrophobic molecules into the outer membrane (9). We have recently described another mechanism of MV biogenesis that involves explosive cell lysis events which produce shattered membrane fragments that rapidly anneal into MVs (10). In *Pseudomonas aeruginosa* explosive cell lysis occurs as a consequence of degradation of peptidoglycan by the muralytic endolysin, Lys, similar to those used by lytic bacteriophage to lyse its bacterial host. Indeed *lys* is encoded in the R-and F-pyocin gene cluster which produces tailocins related to P2 and lambda bacteriophage (11,12).

Bacteriophages (phages) are the most abundant entities on the earth and are commonly found where bacteria exist (13). Bacteriophages are categorised as virulent or temperate based on their life cycle. The lytic life cycle of virulent phage begins when it hijacks the host replication machinery to synthesise phage nucleic acid and proteins, which are then assembled into phage progenies and released from the host after lysing the bacterial cell envelope (14). Temperate phage on the other hand, upon infection may either enter the aforementioned lytic cycle or begin the lysogenic lifecycle whereby the phage DNA is integrated into the host chromosome and replicate as part of its genome as a prophage (15). Prophage integration is not permanent, and under various stress conditions, prophage can excise and enter the lytic lifecycle resulting in host cell lysis.

Phage mediated bacterial lysis is a temporal process, which requires expression of a set of lytic genes, including those encoding endolysins, holins and spanins (11,16,17). The phage endolysin is produced during phage replication and accumulates within the host cytoplasm (16). The translocation of the endolysin through the inner membrane is generally mediated by the holins, which polymerise to form transmembrane holes allowing the phage endolysin to access the cell wall peptidoglycan (11,16). The endolysin mediated peptidoglycan hydrolysis is sufficient to lyse Grampositive bacteria, however an additional membrane spanning protein spanin is thought to be required to complete cell lysis in Gram-negative bacteria (17,18). Phage mediated lysis of Gram-negative bacteria has been reported as a cell bursting phenomenon that appears to be similar to that of tailocin-mediated explosive cell lysis of *P. aeruginosa* (11,18).

Interestingly, MVs are often evident in transmission electron micrographs of bacteriophage preparations (19–21). However, the source of MVs in phage lysates and their association with bacteriophage infection is unclear. In this study we utilised live-cell super-resolution microscopy of *E. coli* cells infected with lytic bacteriophage T4 or T7 and determined that these induced explosive cell lysis events and the production of MVs derived from shattered membrane fragments. We also observed that infection with these bacteriophages was associated with the formation of membrane blebs in intact bacteria. Transmission electron microscopy (TEM) revealed different classes of MVs within phage lysates, consistent with multiple mechanisms of MV biogenesis. To our knowledge this is the first study definitively linking phage infection with any mechanism of MV biogenesis.

## Results

### Live cell imaging of bacterial lysis associated with phage infection

To observe changes within the bacterial cell population due to phage infection we utilised a standard assay for phage propagation in liquid medium (22). *E. coli* strain MG1655 was infected with phages (T4 and T7 suspended in lambda diluent) or a control (lambda diluent) and imaged for 1 h using time-lapse phase-contrast microscopy. The addition of either T4 or T7 phage resulted in lysis of many bacterial cells, whereas no lysis was observed in the control samples (Figure 1A-C, Movie S1-S2). Indeed, in the negative control, bacteria cells were observed to be multiplying, increasing their number within the field of view (Movie S3). Approximately 50% of cell lysis events occurred within 17-32 min for T4 phage-infected cultures and within 8-24 min for T7 phage-infected cultures (Figure 1D). Interestingly, T7 phage lysed more *E. coli* cells compared to T4 phage (approx. 91% and 56%, respectively) over the analysed time period (Figure 1D). These observations indicated that lysis was due to phage infection and not due to any extraneous condition or any experimental artefacts.

**Figure 1:**
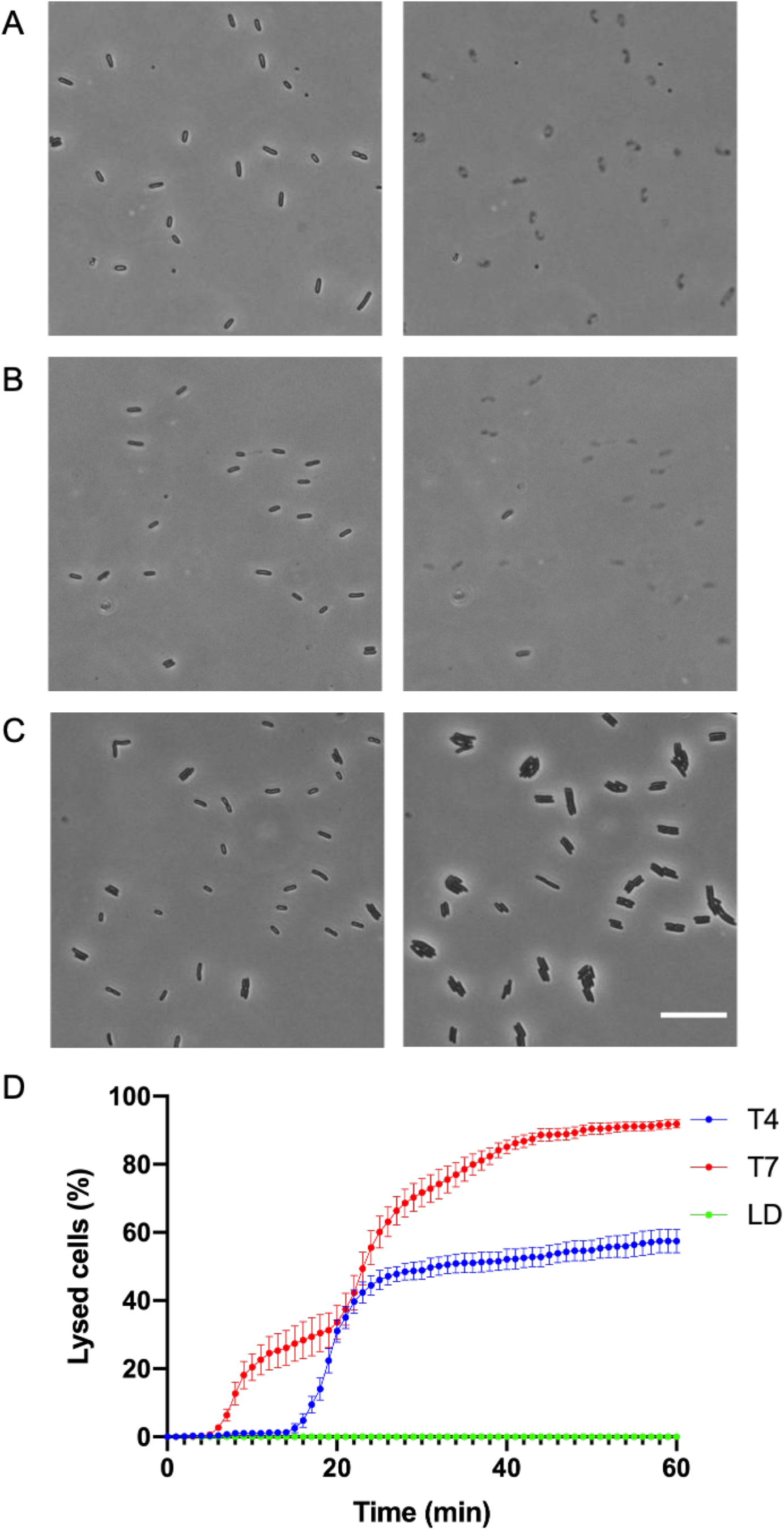
Visualisation of T4 and T7 phage lysing *E. coli*. Time-lapse image sequences of *E. coli* MG1655 treated with (A) T4 phage (B) T7 phage or (C) lambda diluent (negative control). The black thin rods in each frame are live bacteria and the grey, ghosted shapes are lysed cells. Images were taken at 1 frame/min for 1 h at 37°C using phase-contrast microscopy. Scale bar is 20 μm. (D) Percentage of lysed *E. coli* K-12 strain MG1655 cells treated with T4 phage (blue), T7 phage (red) and lambda diluent (LD) control (green) at each 1 min increment for a total of 1 h. The data is presented as mean ± SD (n=3). A total of 1,170 cells were followed for lysis for T4, 1,669 for T7 and 1,470 for lambda diluent. The full time-lapse image series for A-C are presented in Movies S1-S3.

### T4 and T7 phages mediate explosive cell lysis in *E. coli*

To further characterise phage-mediated cell lysis of *E. coli*, single-cell lysis events were visualised using high magnification time-lapse phase-contrast microscopy (Figure 2A-C). This revealed that rapid lysis of rod-shaped cells occurred at either the cell pole (Figure 2A; Movie S4) or mid-cell (Figure 2B; Movie S4). Some cells were also seen to transition from a rod shape to a spherical morphotype prior to rapid cell lysis (Figure 2C; Movie S4). For lysis due to T4 phage infection, the majority of lysis events (64/103; 62%) occurred after expulsion of cellular contents at the cell pole, compared to lysis events due to expulsion of cellular contents at mid cell (18/103; 18%) or after a transition to a spherical morphotype (21/103; 20%) (Figure 2D). For T7 phage infection, the majority of cells lysed after transition to a spherical morphotype (106/118; 90%), with very few cells lysing due to expulsion of cellular contents at the cell pole (6/103; 5%) or mid cell (6/103; 5%). For lysis events that showed the rounding of cells, the most common morphology observed before lysing was a prolate sphere. The phenomenon of these rod-shaped cell *E. coli* cells transitioning into a spherical morphotype prior to cell lysis is similar to the rod to spherical morphotype transition prior to the Lys-mediated lysis of *P. aeruginosa*, which we have previously termed explosive cell lysis (10). Despite some *E. coli* cells lysing without transition to a spherical morphotype, the nature of the lysis is also explosive (Figure 2; Movie S4) and thus suggests that T4 or T7 phage infection can induce explosive cell lysis in *E. coli* from either a rod or spherical morphotype.

**Figure 2:**
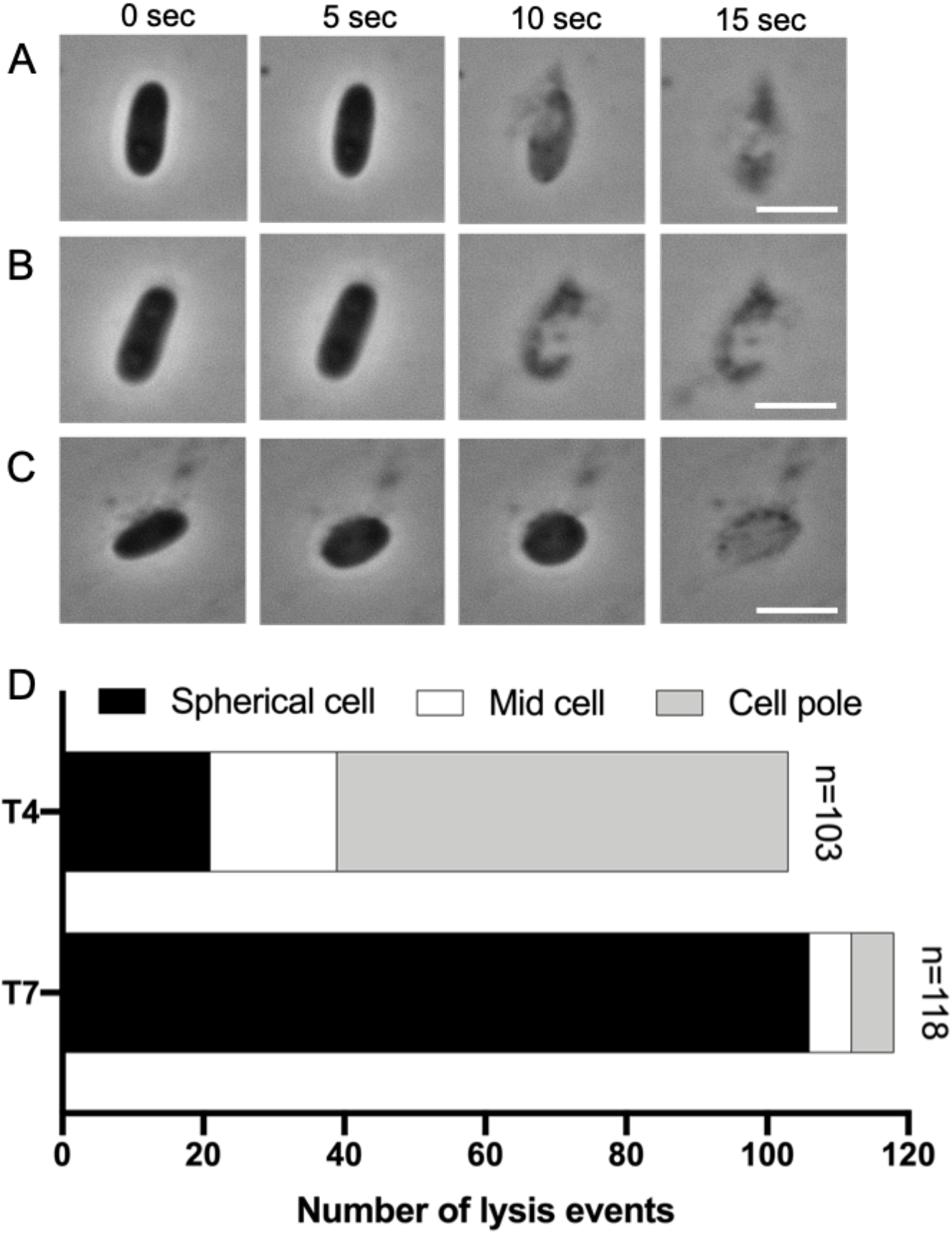
T4 and T7 phage mediate explosive cell lysis in *E. coli*. Time-lapse representative images of *E. coli* MG1655 treated with T4 bacteriophage showing (A) lysis near the cell pole of a rod-shaped cell, (B) lysis near mid cell of a rod-shaped cell or (C) lysis of a cell that has transitioned from a rod to a spherical morphotype. Images were taken using phase contrast and are representative of at least 4 independent experiments. Scale bar is 2 μm. (D) Total number of events separated according to where in the *E. coli* cell lysis occurred, or the cell morphotype prior to lysis, when infected with T4 or T7 phage. The full time-lapse image series for A-B are presented in Movie S4.

### T4 and T7 phage-mediated explosive cell lysis of *E. coli* results in MV biogenesis

We have previously reported that explosive cell lysis in *P. aeruginosa* results in MV biogenesis (10). We therefore explored the possibility that lytic phages could be responsible for MV formation through a similar mechanism. To examine this, we followed multiple cell lysis events that resulted from all 3 lysis patterns (Figure 2) of *E. coli* infected with T4 or T7 phage using 3D-structured illumination microscopy (3D-SIM) and the lipophilic dye FM1-43 to visualise cell membranes and MVs. We observed numerous MV-like structures within regions of extensive cell lysis (Figure 3A), whereas no such MVs were observed on control samples, where there was no lysis (Figure 3B).

**Figure 3:**
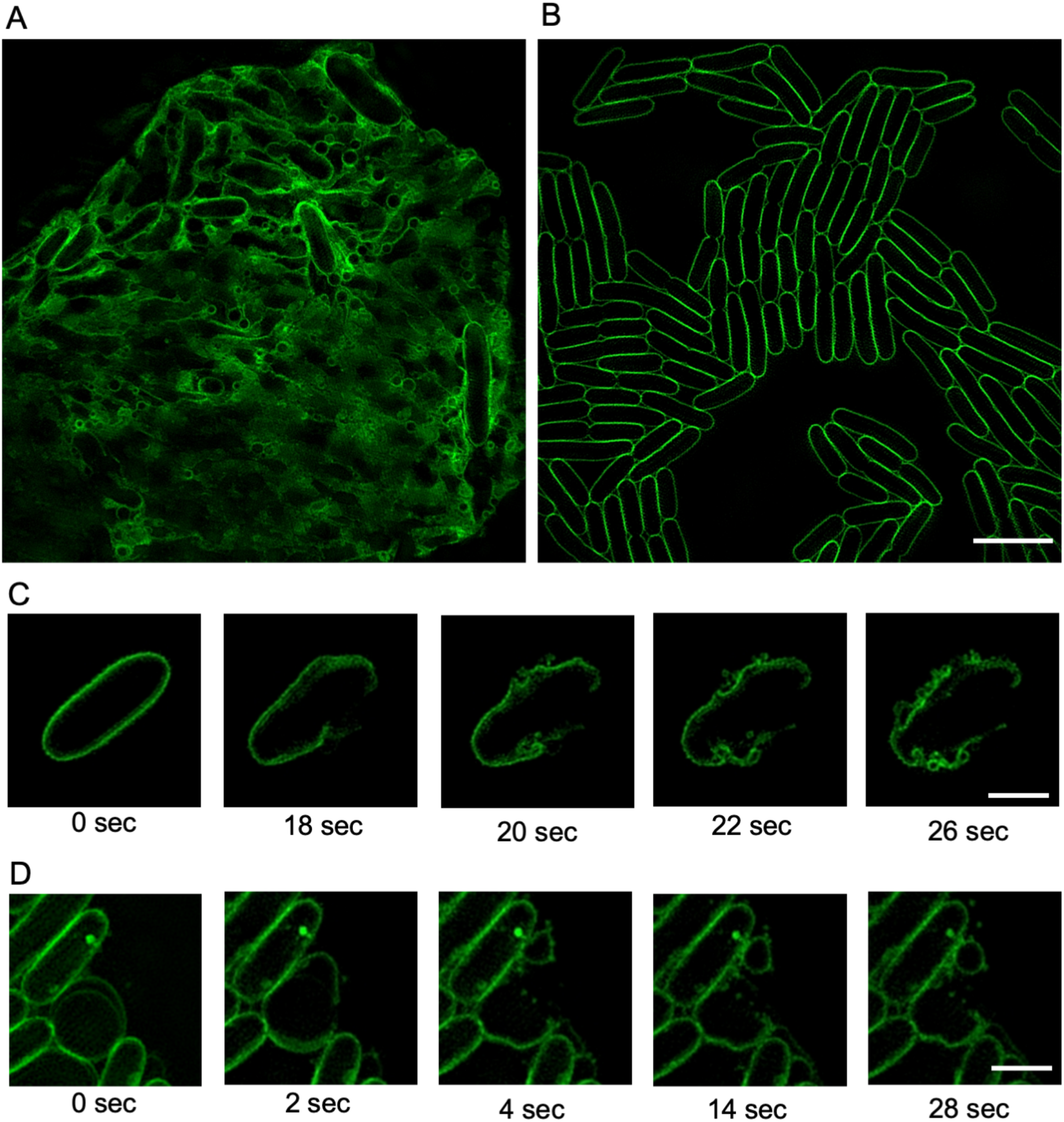
Explosive cell lysis generates MVs in *E. coli*. (A-B) 3D-SIM images of *E. coli* MG1655 (A) infected with phage T7 or (B) treated with lambda diluent control. Images of *E. coli* infected with T4 phage were similar to A. (C-D) Time-lapse image sequences of explosive cell lysis events of *E. coli* MG1655 cells infected with (C) T4 or (D) T7. The membrane dye FM1-43X (green) was used for membrane staining. Images were acquired using 3D-SIM and are representative of 67 (T4) or 57 (T7) fields of view imaged over 3 separate days. Scale bar is 5 μm (A-B) or 2 μm (C-D). The full time-lapse image series for C-D are presented in Movies S5-S6.

We used live 3D-SIM to follow the fate of cell membranes. We found that explosive cell lysis events produced membrane fragments that quickly vesicularised to form circular MVs that persisted in the same topology thereafter (Figure 3C-D; Movie S5-S6). Interestingly, in some instances we could see clear separation of the IM and OM in cells prior to lysis (Figure 3D, Movie S6). MVs derived from explosive cell lysis events were found to form regardless of the pattern of individual cell lysis (originating from the cell pole, mid-cell or spherical cell), which suggests that phage-mediated explosive cell lysis could be a major route of MV biogenesis.

### T4 and T7 phage infection also results in MV formation through membrane blebbing

In addition to the MVs produced after cell lysis, a large number of membrane blebs were also observed in bacterial populations that were not lysed under phage-infected conditions. Since in Gram-negative bacteria blebbing and explosive cell lysis have been suggested as the two different routes for the MV biogenesis (9), we utilized live cell 3D-SIM to investigate whether phage infection could also trigger MV formation through blebbing in *E. coli*. Strikingly, we observed multiple MVs blebbing from many cells when infected with either T4 or T7 phage (Figure 4; Movie S7-8), and no MV blebs visible on cells treated with lambda diluent (control). This demonstrates that MV blebbing from intact cells was due to phage infection and not an artefact of fluorescent imaging, the addition of FM1-43 dye or other factors.

**Figure 4:**
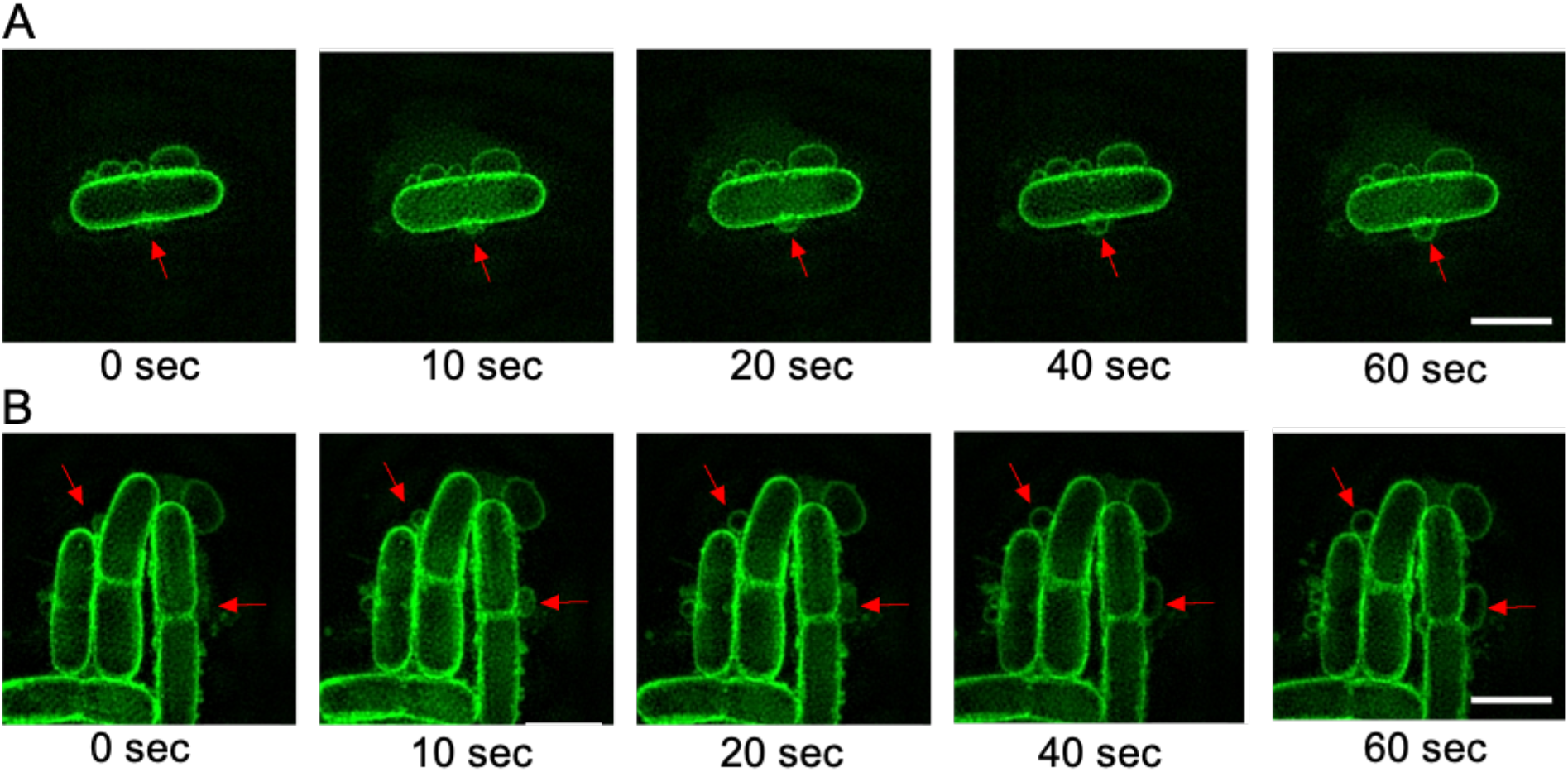
Phage infection induces MV blebbing in the absence of cell lysis. Time-lapse image sequence of membrane blebbing in *E. coli* MG1655 treated with (A) T4 phage or (B) T7 phage. Red arrows indicate site of MV blebs on cell. The membrane dye FM1-43X (green) was used for membrane staining. The lambda diluent control in this experiment was visually identical to Figure 3B. Images were acquired using 3D-SIM from two (T4) or four (T7) independent experiments and are representative of 28 (T4) or 99 (T7) cells with MV blebs. Scale bar is 2 μm. The full time-lapse image series for A-B are presented in Movies S7-S8.

### Phage infection results in different types of MVs in phage lysates

Our observations suggest that there are multiple routes by which MVs are produced as a consequence of phage infection: membrane blebbing from intact cells that gives rise to OMVs; and explosive cell lysis events which can lead to the formation of MVs comprised of both outer and inner membranes (OIMVs) as well as explosive outer membrane vesicles (EOMVs) (9).

Due to the resolution limitation of 3D-SIM we were unable to obtain detailed structures of the MVs. We therefore performed Transmission Electron Microscopy (TEM) of phage lysates to examine the morphology of MVs in lysates obtained from phage infected cultures. As expected, under TEM, both T4 and T7 phage lysates showed the respective phage, MVs and cellular debris (Figure 5A and 5B). Variable MV sizes were observed and overall T7 infected cells produced significantly larger MVs than T4 (180.9 nm ± 48.0 compared to 126.8 nm ± 53.9 for T7 or T4, respectively) (Figure 5D). We also observed two general morphotypes of MVs within both T4 and T7 lysates (Figure 5C), which we described as: (i) OMVs consisting of a single membrane layer or (ii) OIMVs consist of both inner and outer membrane layers. TEM was unable to differentiate between OMVs derived by blebbing or explosive cell lysis.

**Figure 5:**
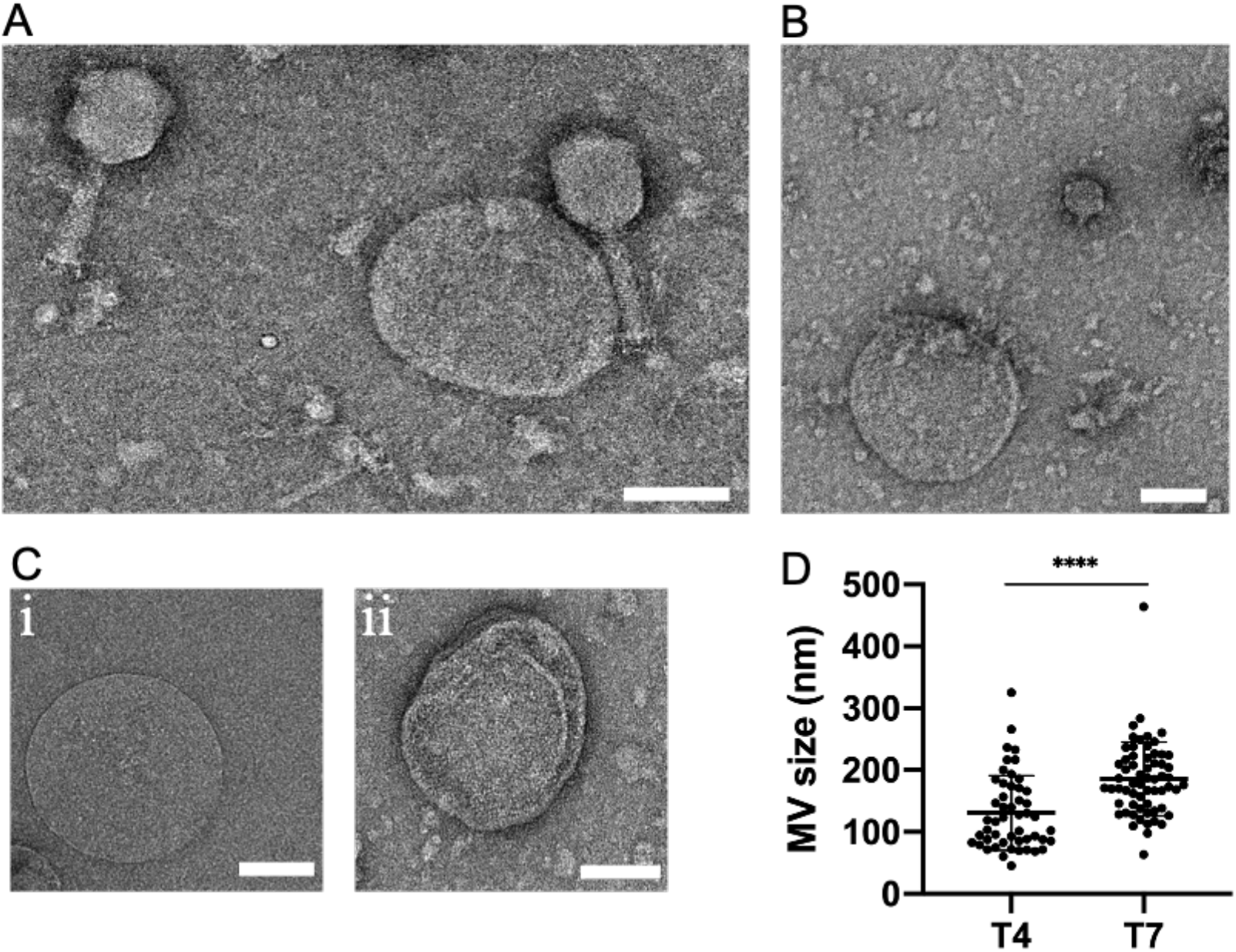
Phage infection of *E. coli* produces a range of different kinds of MVs within phage lysates. Representative TEM images of (A) T4 lysate with MV and cell debris, (B) T7 lysate with MVs and cell debris. (C) Representative images of different forms of MVs observed within both T4 and T7 lysates: (i) classic OMV or (ii) OIMV. (D) MVs sizes within T4 (50 MVs analysed) and T7 (62 MVs analysed) lysates. Data are presented as the mean ± SD (\*\*\*\**P*<.0001, unpaired t-test with Welch’s correction). For A and B images are representative of 49 (T4 lysate) or 61 (T7 lysate) random fields of view imaged in 3 independent experiments. Scale bar is 100 nm.

## Discussion

In this study, we have demonstrated that T4 and T7 phage infection of *E. coli* generates MV production via both explosive cell lysis and membrane blebbing. Using 3D-SIM we observed a range of sizes of MVs in phage infected cultures, which was also observed in phage lysates using TEM. Notably, we observed MVs from both the T4 and T7 phage to be within the range of 50 – 300 nm, which correlates with the sizes of bacterial MVs reported previously (9,10,25,26).

In the case of explosive cell lysis we observed three main patterns of lysis: where cells remained as rods and lysed at either the cell pole or mid cell or where rods transitioned to a spherical morphotype before lysing. Interestingly, we found that the majority of lysis with T4 phage infection occurred at the poles of rods, whereas with T7 the majority of lysis occurred after a cell transitioned to a spherical morphotype. Both patterns of phage-mediated lysis have been previously reported to be associated with phage holin-endolysin systems (18). Berry *et al*. (2012) demonstrated that infection of *E. coli* MC4100 with lambda phage consisting of a canonical holin-endolysin system mainly caused lysis of rod-shaped cells. In contrast they demonstrated that an isogenic hybrid phage consisting of a pinholin-SAR endolysin mainly caused lysis after transition of cells to a spherical morphotype (18). Strikingly, in the current study we found that despite both T4 and T7 phage both possessing a canonical holin-endolysin system (11,16), these phages had two distinct sets of lysis patterns. This finding suggests that phage mediated cell lysis may not be solely dependent upon which holin-endolysin system the phage possesses but could also depend on other factors. For instance, the T4 phage possesses an endolysin with muramidase activity (PF00959) that acts on the glycosidic bond (between N-acetylmuramic acid and N-acetylglucosamine) linking the amino sugar in the peptidoglycan (23), whereas the T7 endolysin has zinc amidase activity (PF01510) that acts on the amide bonds between N-acetylmuramic acid and L-alanine of the cross-linking oligopeptide stems and interpeptide bridge in the peptidoglycan (24). Thus, differences in enzymatic activity of individual phage endolysins could be responsible for the observed differences in lysis patterns. One other possibility is that in some cells the holin and/or endolysin may accumulate at different sites or in different amounts within the cell envelope resulting in the observed differences in lysis patterns.

We also found that irrespective of the lysis pattern, lysis events were explosive and produced MVs from the shattered membrane fragments. These observations are in agreement with our previous observations of explosive cell lysis in *P. aeruginosa* where we observed MVs forming through curling and self-annealing of shattered membrane fragments from lysed cells (10). MV biogenesis via explosive cell lysis is mediated by holin-endolysin systems in *P. aeruginosa* (10, 27) and is likely to also be occurring as a consequence of the holin-endolysin systems of lytic T4 and T7 phage in *E. coli*. This process likely accounts for MV formation in other Gram-negative bacteria.

Additionally, we found that phage infection not only produces MVs through cell lysis but that it also triggers MV formation through a membrane blebbing mechanism from intact cells. It is currently unknown what the mechanisms are that direct phage infected cells to undergo explosive cell lysis or to remain intact and generate MVs via membrane blebbing, however this would be interesting to investigate in future work.

Collectively, our findings revealed that *E. coli* infected with lytic phage produce MVs through both explosive cell lysis as well as membrane blebbing. Given that bacteriophages are the most abundant entities on earth and infect a wide range of bacteria, our observations suggest that phage mediated MV formation in Gram-negative bacteria may be a major contributory process to the abundance of MVs in nature.

## Materials and methods

### Bacterial strains and bacteriophages

The bacterial strain used in this study was *Escherichia coli* K-12 MG1655 (28). The bacteriophage used in this study were T4 phage which is a Virulent Myovirus that infects *E. coli* (DSM4505) and T7 phage which is a Virulent Podovirus that infects *E. coli* (DSM4623). These phages were obtained from the Leibniz-Institut DSMZ - Deutsche Sammlung von Mikroorganismen und Zellkulturen GmbH.

Bacterial strains used in this study were grown in LB (10 g L^-1^ Tryptone (Oxoid: LP0042), 10 g L^-1^ NaCl (Oxoid: LP0005) and 5 g L^-1^ Yeast extract (Oxoid: LP0021)) as either liquid culture (with shaking at 100 rpm) or solid culture (supplemented with 15 g L^-1^ agar (Oxoid: LP0011B)) at 37°C. Phages were suspended in lambda diluent (10 mM Tris.HCl pH 7.5, 8 mM MgSO_4_).

### Phage lysate preparation and titration

Phage lysate was prepared as per standard procedure described elsewhere (22). Briefly, liquid phage suspension was centrifuged at 4000 *g* for 10 min at 4°C and the supernatant was subsequently filter sterilised using a 0.45 μm pore sized membrane filter (Sartorius Minisart®, 16555). The filtrate phage lysate was stored at 4°C. Phage titre of harvested lysate was determined using spot tests (29).

### Bacterial lysis assay

A liquid culture assay used for phage propagation (22) was adapted to demonstrate phage mediated bacterial lysis (bacterial lysis assay). Briefly, 1 mL overnight culture of *E. coli* strain MG1655 was diluted in 10 mL of fresh LB and incubated at 37°C for 1 h (early log phase). 10 μL of purified phage sample with a titre more than 10^8^ pfu ml^-1^ was then added to the bacterial suspension. 1 μL of the mixture just after phage addition was used for microscopic examination to visualise bacterial lysis and the mixture was incubated for a further 5 h at 37°C with shaking at 100 rpm for phage propagation. A control test with phage diluent instead of phage sample was used to validate the assay.

### Phase-contrast and super-resolution microscopy

For both the phase contrast and super-resolution microscopy cells were applied to gellan gum solidified nutrient media as described previously (30). Briefly, cells from a liquid culture were spotted on nutrient media (10 g L^-1^ Tryptone (Oxoid: LP0042), 10 g L^-1^ NaCl (Oxoid: LP0005), 5 g L^-1^ Yeast extract (Oxoid: LP0021)) solidified with 8 g L^-1^ gellan gum (MP Biomedicals: 180106). The LB gellan gum (LBGG) nutrient media was supplemented with the fluorescent dye FM1-43FX (Life Technologies) to a final concentration of 5 μg mL^-1^ to visualise the cell membrane for super-resolution microscopy. Phase-contrast microscopy was performed using an Olympus IX71 inverted microscope fitted with an environmental chamber (Solent Scientific, Segensworth, UK) and AnalySIS Research acquisition software (Olympus Australia, Notting Hill, VIC, Australia). Super-resolution microscopy was performed using the Delta Vision OMX SR microscope using 3D-SIM mode. Live images were captured using a 1.42 numerical aperture 60x oil objective, standard filter set, a scientific CMOS 512×512 pixel 15-bit camera (PCO AG, Kelheim, Germany) and AquireSR software. Raw 3D-SIM images were obtained through section using a 125-nm Z-step size, which were then reconstructed using SoftWorX software (Applied Precision, GE Healthcare).

### Transmission electron microscopy

For transmission electron microscopy, phage lysates (generated as described above) were deposited on carbon coated Formvar films on copper grids, with an adherence time of 1 min, stained with 2% Uranyl acetate, and examined in a FEI Tecnai electron microscope (Tecnai, T20) at 200 kV acceleration voltage. Images were produced using Gatan camera.

### Image analysis

Phase-contrast microscopy images and movies were analysed using Fiji (v. 2.0.0) (31). Reconstructed 3D-SIM images from super-resolution microscopy were rendered and presented using IMARIS (v. 9.2.1, Bitplane Scientific). Linear adjustments to signal contrast and brightness were made in the images presented but no gamma settings were changed. Transmission electron microscopy images were also analysed using Fiji (v. 2.0.0) (31) and dimension of images were measured with their respective image scale.

### Statistical analyses

Statistical analyses were performed using GraphPad Prism (v. 8.4.3) for Mac OS X, GraphPad Software, San Diego, California USA, www.graphpad.com.

## Acknowledgements

We thank the Microbial Imaging Facility (MIF), UTS for providing access to optical microscopes and the Microstructural Analysis Unit (MAU), UTS for access to the transmission electron microscope.

## Contributions

PKM, NKP and CBW conceived the study and designed the research. PKM and GB performed the experiments. PKM, GB, LMN and CBW analysed results. PKM, GB, LMN, NKP and CBW wrote the manuscript.

## Supplementary movies

**Movie S1: T4 phage lyses *E. coli* MG1655 cells.** *E. coli* was infected with T4 phage cultured on nutrient media. Phase-contrast microscopy was used to image cell lysis with 1 frame/min captured for a total time of 1 h. Movie is played back at 600x real time. Scale bar is 50 μm.

**Movie S2: T7 phage lyses *E. coli* MG1655 cells.** *E. coli* was infected with T7 phage cultured on nutrient media. Phase-contrast microscopy was used to image cell lysis with 1 frame/min captured for a total time of 1 h. Movie is played back at 600x real time. Scale bar is 50 μm.

**Movie S3: Control for Movies S1-S2.** *E. coli* MG1655 treated with lambda diluent cultured on nutrient media. Phase-contrast microscopy was used to image cells with 1 frame/min captured for a total time of 1 h. Movie is played back at 600x real time. Scale bar is 50 μm.

**Movie S4**: **Phage infection results in explosive cell lysis**. *E. coli* MG1655 was infected with T4 phage and cultured on nutrient media. Imaging was performed with phase-contrast microscopy captured at 1 frame/5 sec for a total time of 500 sec. Movie is played back at 50x real time. Scale bar is 5 μm.

**Movie S5: T4 phage-mediated explosive cell lysis produces MVs.** *E. coli* MG1655 was infected with T4 phage and cultured on nutrient media containing the membrane stain FM1-43X. Imaging was performed with 3D-SIM captured at 1 frame/2 s for a total time of 170 sec. Movie is played back at 20x real time. Scale bar is 1 μm.

**Movie S6: T7 phage-mediated explosive cell lysis produces MVs.** *E. coli* MG1655 was infected with T7 phage and cultured on nutrient media containing the membrane stain FM1-43X. Imaging was performed with 3D-SIM captured at 1 frame/2 s for a total time of 170 sec. Movie is played back at 20x real time. Scale bar is 1 μm.

**Movie S7: T4 phage infection induces membrane blebbing.** *E. coli* MG1655 was infected with T4 phage and cultured on nutrient media containing the membrane stain FM1-43X. Imaging was performed with 3D-SIM captured at 1 frame/2 s for a total time of 170 sec. Movie is played back at 20x real time. Scale bar is 1 μm.

**Movie S8: T7 phage infection induces membrane blebbing.** *E. coli* MG1655 was infected with T7 phage and cultured on nutrient media containing the membrane stain FM1-43X. Imaging was performed with 3D-SIM captured at 1 frame/2 s for a total time of 170 sec. Movie is played back at 20x real time. Scale bar is 1 μm.

